# Assessing the quality of supplementary sensory feedback using a crossmodal congruency task

**DOI:** 10.1101/194977

**Authors:** Daniel Blustein, Adam Wilson, Jon Sensinger

## Abstract

Peripheral nerve interfaces show promise in making prosthetic limbs more biomimetic and ultimately more intuitive and useful for patients. However, approaches to assess these emerging technologies are limited in their scope and the insight they provide. When outfitting a prosthesis with a new feedback system it would be helpful to quantify its physiological correspondence, i.e. how well the experimental feedback mimics the perceived feedback in an intact limb. Here we present an approach to quantify physiological correspondence using a modified crossmodal congruency task. We trained 60 able-bodied subjects to control a bypass prosthesis under different feedback conditions and training durations. We find that the crossmodal congruency effect (CCE) score is sensitive to changes in feedback modality (multi-way ANOVA; F(2,48) = 6.02, p<0.05). After extended training, the CCE score increased as the spatial separation between expected and perceived feedback decreased (unpaired t-test, p<0.05). We present a model that can quantitatively estimate physiological correspondence given the CCE result and the measured spatial separation of the feedback. This quantification approach gives researchers a tool to assess an aspect of emerging augmented feedback systems that is not measurable with current motor assessments.

## Introduction

The performance of clinically-available upper-limb prostheses has been partly hindered by a lack of intuitive and useful feedback^1,2^. Direct neural stimulation to convey force or other feedback to a user controlling a prosthetic hand may lead to improved systems that better mimic the dynamics of the intact human hand. These peripheral nerve interfaces (PNIs) with bidirectional communication between device and body have been shown effective in controlled settings^3-5^. Efforts to improve long term viability, once a concern for such invasive interfaces, have also advanced using wireless signal transmission^6^, osseointegration approaches^7^, and stable electrode designs^8^. While the promise of PNIs is apparent, the development of feedback assessment tools has lagged these emerging technologies.

Traditional performance-based movement assessments may not capture the overall quality of a novel feedback modality. Standard clinical motor assessments such as the Box and Blocks Test^9^, the Nine Hole Peg Test^10^, the Southampton Hand Assessment Procedure^11^ and the Assessment of Capacity for Myoelectric Control^12^ focus on quantifying motor performance but do not provide insight into other potential factors of particular feedback systems. Emerging feedback systems may provide intrinsic value beyond motor performance gains. Evidence suggests that user-trusted feedback can lead to aspects of incorporation and embodiment^13^, that could improve prosthesis acceptance^2^, reduce phantom pain^14,15^ or provide other benefits that would not be detected by traditional motor assessments^16^.

Emerging PNI studies have relied on qualitative subjective user descriptions of feedback quality^4,17^. Users report the location and sensation of the applied experimental feedback and the quality of that feedback is inferred. For example, in one study subjects described sensations in terms of pressure, tingle, vibration, or light moving touch^4^. As stimulation intensity was varied, one subject described the sensation as changing from “tingly” to “as natural as can be”^4^. Some conclusions are clearly justifiable, for example, self-described “natural, non-tingling feedback” is presumably better than “uncomfortable, deep, dull vibration”^4^. But for more complete comparisons the subjective approach lacks consistency and reliability. When a user perceives touch through a PNI, how closely does that percept match the touch sensation perceived with intact anatomy? Experimental feedback with low physiological correspondence may be difficult to interpret, or even painful^18^. In this study we have sought to quantify the physiological correspondence of supplementary sensory feedback modalities.

While quantifying the physiological correspondence of an experimental feedback system is the ultimate goal, PNI feedback can also be spatially misaligned from what it strives to mimic. The perceived location of the experimental feedback may differ from the expected feedback location, which may greatly affect the feedback’s usability. For example, a visually-observed contact on a prosthetic fingertip may be felt, as the result of neural stimulation, several centimeters away at the base of the finger (see Fig. 1, top right panel)^17^. Quantifying physiological correspondence must also consider the effect of misaligned feedback percepts.

**Figure 1.**
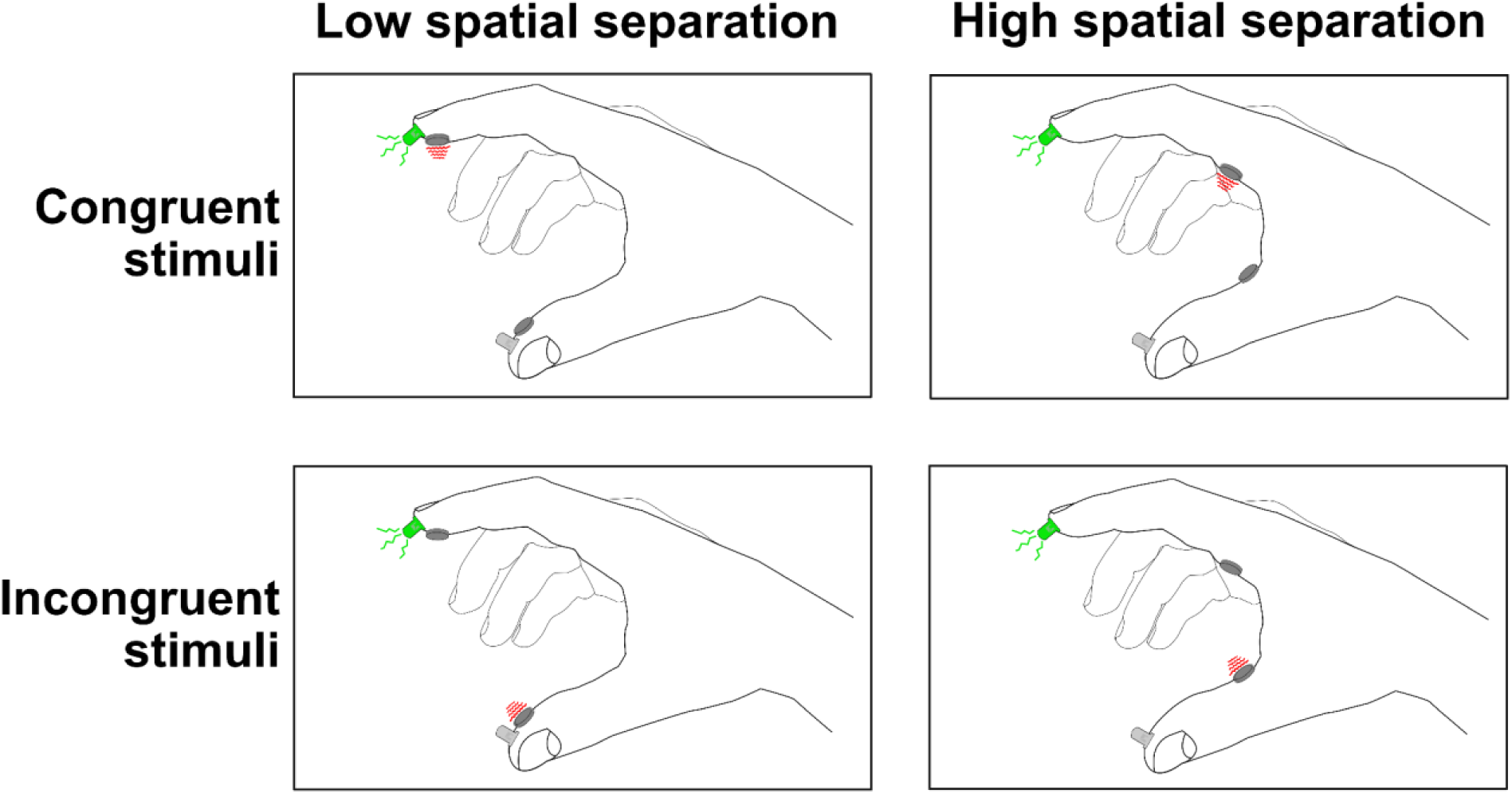
Crossmodal congruency effect framework. Subjects rapidly select the experimental feedback (e.g. vibration shown in red). The crossmodal congruency effect (CCE) score is the difference between the response time to congruent and incongruent stimuli. Spatial separation is the physical distance between the two paired sensory percepts. Incongruent stimuli are defined as unpaired stimuli presented simultaneously (bottom row).

The crossmodal congruency effect (CCE) score provides an objective measure of incorporation of a feedback modality^19,20^. The degree of feedback incorporation is affected by the spatial separation and physiological correspondence of the experimental feedback^19,21,22^ (Fig. 1). Given that a person’s CCE score and spatial separation for a particular feedback modality are directly measurable, physiological correspondence can be quantified using an explanatory model.

To develop a model that can quantify physiological correspondence of experimental feedback, we completed a comprehensive study of 60 able-bodied subjects controlling a bypass prosthesis under different feedback conditions. We found that incorporation as measured by CCE score changes with feedback modality. After extended training, CCE score increased as the spatial separation between expected and perceived feedback decreased. From the results we developed a model to estimate the physiological correspondence of a feedback modality given a CCE score and a measured spatial separation. The approach we present provides an important first step towards quantitatively measuring the degree to which feedback actually feels physiologically accurate.

## Results

To determine if experimental feedback provided to a person could be incorporated, we measured the CCE score^19,23^ of 60 able-bodied individuals after training with a bypass prosthesis^24^(Fig. 2). Participants trained using the bypass prosthesis to move mechanical eggs with one of three feedback modalities: vibration, electrical stimulation or skin deformation^24^. Training duration (short vs. extended) and spatial separation between the expected feedback on the fingertip contact point and the perceived feedback (matched on fingertip vs. >12 cm away) were also varied (see Supplementary Table S1 for all experimental conditions). We report the sensitivity of the CCE score to feedback modality and spatial separation. We use the results to generate a model that can estimate the physiological correspondence of a feedback type.

**Figure 2.**
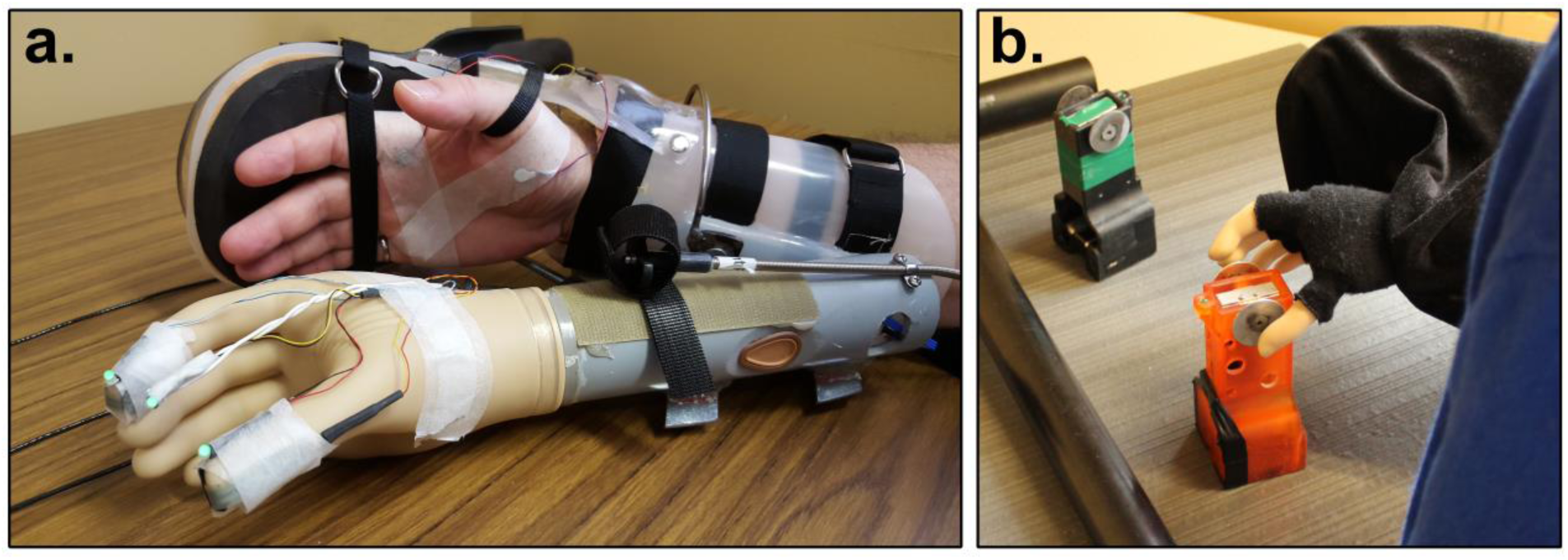
Experimental setup. **a.** Bypass prosthesis to allow able-bodied subjects to control a prosthesis with modifiable feedback. The sensorized prosthesis could detect force applied to the thumb and the index finger and provide proportional feedback to the user conveyed as vibration, electrical stimulation or skin deformation. The attached LEDs are used during the CCE score measurement protocol (see Methods). During training and testing the intact hand and harness are covered with black fabric. **b.** Variable weight mechanical eggs were moved during training periods. Load cells on eggs detected grasp force and simulated an egg breakage with a light cue when a threshold was exceeded.

We observed a main effect of feedback modality on CCE score (Fig. 3). A three-way ANOVA with CCE score as the independent variable and three categorical dependent variables representing spatial separation, training level and the three feedback modalities resulted in a statistically significant effect of feedback (F(2,48) = 6.015, p<0.01, ω=0.28). Interactions terms between independent variables were not significant and not included in further analysis (p>0.05). Bonferroni post-hoc tests showed that the CCE score was significantly higher for vibration feedback [μ=120.49ms ± 53.25] compared to skin deformation feedback [μ=70.99ms ± 46.04] (p<0.05) and non-significantly higher than electrical stimulation [μ=84.06ms ± 35.5] (p=0.052). A higher CCE score indicates a higher level of incorporation for that feedback modality^19^. The trends observed with data binned for each modality are also seen with the CCE scores of each of the 12 treatment groups (Supplementary Fig. S1). Changes in the provided feedback modality had a significant effect on the measured CCE score.

**Figure 3.**
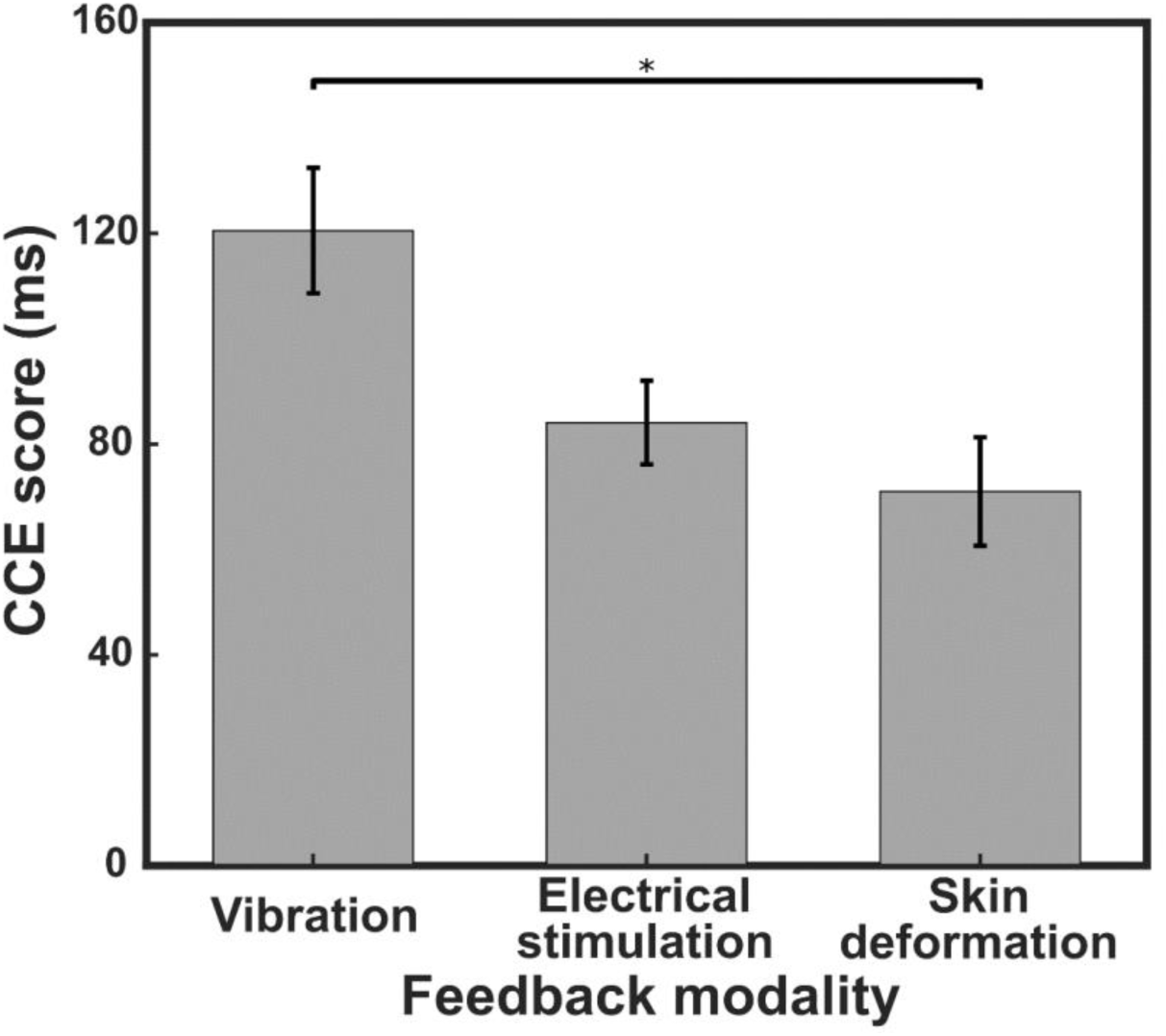
Feedback modality affects CCE score. Means and standard error plotted for 20 subjects in each feedback modality group. Statistical significance verified using three-way ANOVA with Bonferroni post-hoc comparison. * = p<0.05.

The spatial separation between perceived and expected feedback affects CCE score, but only after extended training (Fig. 4). Due to the global differences in CCE scores across modality (see Fig. 3), we use CCE scores normalized within each modality to observe the effect of spatial separation. In subjects with long training periods, there was a significant effect of spatial separation on normalized CCE score (unpaired t-test, p<0.05)(Fig 4a). CCE scores decreased as spatial separation increased, indicating that incorporation diminished as the perceived feedback did not align with the expected feedback location. Spatial separation appeared to have no effect on CCE score in subjects with a short training period (unpaired t-test, p>0.05)(Fig 4B). Short training involved 50 minutes of practice moving mechanical eggs and extended-training lasted 80 minutes. Since spatial separation only had a significant effect on the measured CCE score after extended training, we do not include the short training data in further analysis.

**Figure 4.**
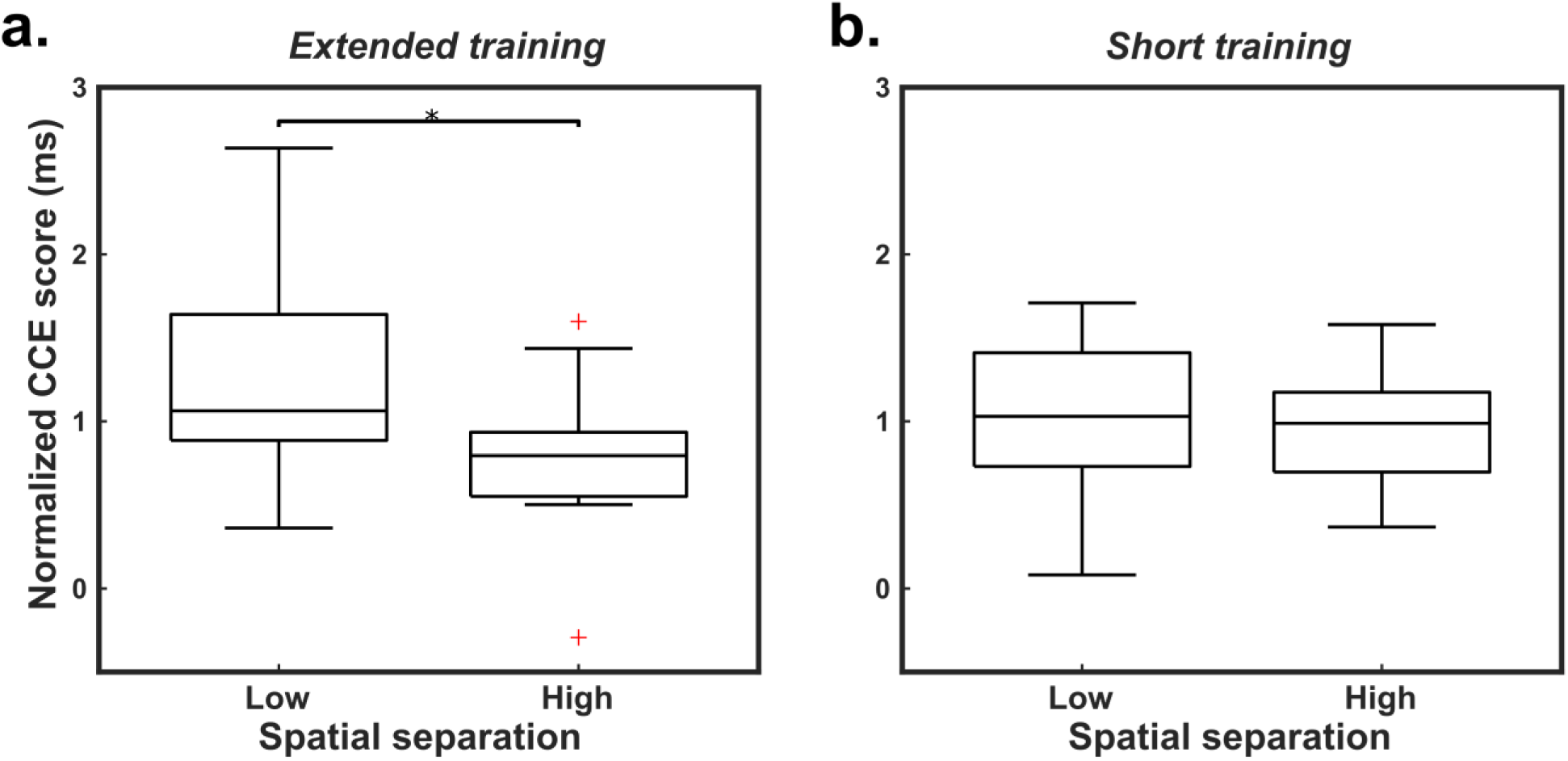
Spatial separation affects CCE score after training. **a.** CCE score decreased with increased spatial separation for extended-training subjects. **b.** There was no observed effect of spatial separation on CCE score after short periods of training. Statistical significance tested using unpaired two-sample equal variance t-tests: * = p<0.05. Reported results are normalized to the group mean within each feedback type to account for global differences across modalities.

Given the significant effects of feedback modality and spatial separation after extended training on CCE score, we attempted to fit a model to estimate the physiological correspondence of the provided feedback. Feedback modalities differ in their level of physiological correspondence to intact biological feedback. Thus we defined physiological correspondence numerically as the average CCE score for all treatment groups within a modality (i.e. the normalization factors used in Fig. 4). This allowed for the conversion of the feedback modality categorical variable into a continuous scale. Since CCE score is a function of physiological correspondence and spatial separation^19,21,22^, we fit a multiple linear regression to the data from the 30 extended-training subjects (F(2,27) = 4.93, p < 0.05, R^2^=0.27). The model can be expressed as

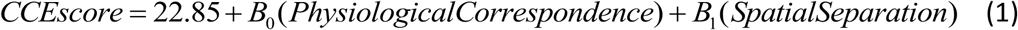

where B_0_ = 0.98 (p < 0.05) and B_1_ = -37.6 (p < 0.05). The spatial separation term was defined as zero for spatial separations of 3.0cm or less, and one for spatial separations greater than 12.0cm. Given measured values for CCE score and spatial separation, the physiological correspondence of a feedback system can be estimated with this model as

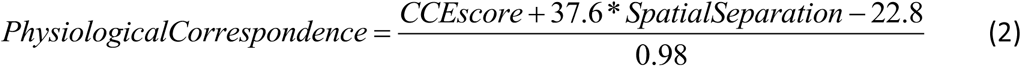

This equation was used to calculate the physiological correspondence for all subjects. The effect of spatial separation on CCE score (Fig. 4a) was observed as trends within the extended-training results for each modality (Fig. 5, top panel). The effect of spatial separation is not present when observing physiological correspondence results converted from the CCE scores of the same subjects (Fig. 5, bottom panel), supporting the model’s ability to account for the effect of spatial separation.

**Figure 5.**
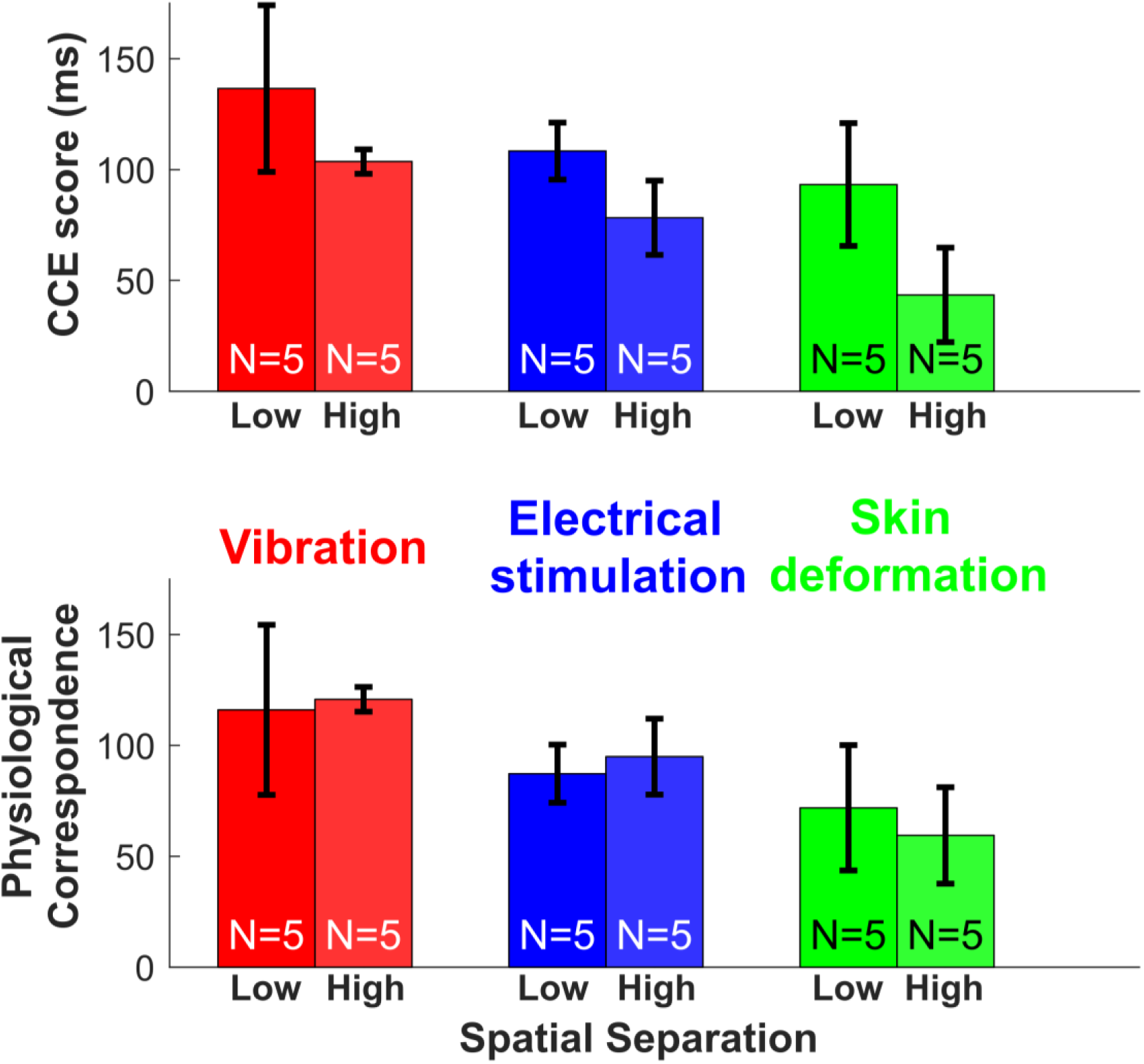
Physiological correspondence quantification is not affected by spatial separation. **Top panel.** CCE score means and standard error for the extended-training subjects. For each modality, the CCE score is lower for the high spatial separation group. **Bottom panel.** Physiological correspondence results for the extended-training subjects. The effect of the spatial separation observed in the CCE score results is not present.

We further analyzed the data to investigate possible explanations for the low CCE score observed with skin deformation feedback. The low CCE score could not be attributed to the latency of the skin deformation feedback application (Supplementary Data S1). Additionally, we observed a non-significant inversely proportional trend between motor performance (movement success rate during training) and CCE score (Supplementary Fig. S2).

## Discussion

We demonstrated that CCE scores can be used to assess feedback quality, specifically the physiological correspondence of a feedback modality. We collected data from 60 able-bodied subjects controlling a bypass prosthesis and developed a predictive model to output physiological correspondence estimates. This model can be applied to other novel sensory feedback systems, such as amputees using peripheral nerve interfaces. The psychophysics-based technique we have presented fills a need for more informative assessment of advanced prosthetic systems.

Given the physiological correspondence estimation model (equation (2)) and the extended-training data we collected, we provide a scale to contextualize physiological correspondence scores (Fig. 6). Researchers who measure the CCE score and spatial separation of a feedback system can calculate the physiological correspondence using equation (2) and compare the results to the scale presented to better classify their conclusions. For example, if a feedback modality’s assessed physiological correspondence is 130, then the feedback would have a similar level of correspondence to vibration feedback. This example feedback modality would have a high level of physiological correspondence. Researchers assessing the physiological correspondence of a novel feedback modalities can use the scale in Fig. 6 as a benchmark for the analysis of results.

**Figure 6.**
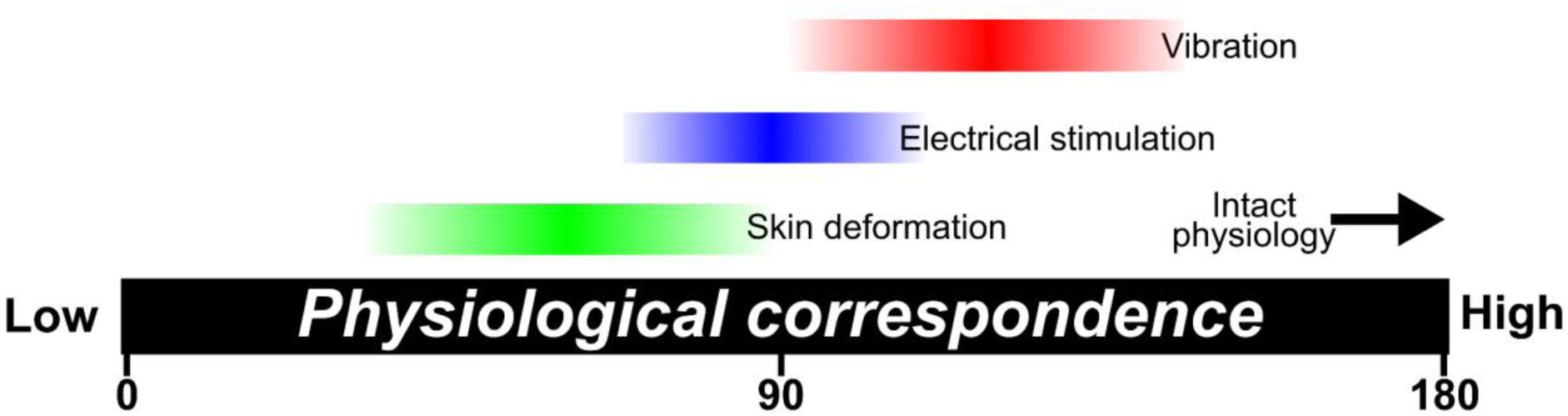
Physiological correspondence benchmark scale. Benchmark data for different feedback modalities to allow for comparison and contextualization of results from the assessment of novel feedback systems.

Our results support previous findings^19,25^ that CCE score is influenced by feedback modality and spatial separation, i.e. the offset between visually-observed feedback and the location of experimentally-provided feedback. Although PNIs strive to minimize spatial separation, the imprecision of neural stimulation makes the generation of perfectly-aligned percepts quite difficult. A high spatial separation will affect the incorporation of the feedback, resulting in a lower CCE score^19^. In regard to different types of feedback, the dynamics and timing of experimental feedback can affect its subjective user-assessment of “naturalness”^4^. Researchers often strive to elicit natural feeling percepts with experimental feedback systems. But to our knowledge we have not seen any attempts at objectively quantifying this “naturalness” sensation. We use physiological correspondence, i.e. how well a feedback modality mimics the feedback experienced with intact anatomy, as a proxy for “naturalness”. Since CCE score and spatial separation are easily measurable, we can use the relationship between these three variables to quantify the physiological correspondence of supplementary sensory feedback.

One potential limitation of the model we developed is that CCE scores may be affected by additional factors besides physiological correspondence, spatial separation and training. We did not factor in the effect of fatigue, time of day, baseline reaction time, and feedback characteristics such as latency, consistency and dynamics. Variability in feedback characteristics were limited by our use of a real-time embedded system to provide feedback with correspondingly low latency (<30ms for vibration and skin deformation). However, the method used to calibrate the skin deformation feedback may have led to variable levels of incorporation. For example, the tactor starting position was not clearly visible in the experimental setup and for some subjects it may have been in contact with the skin or arm hair at zero force levels. Although we could not eliminate the effects of all potential factors affecting CCE score, physiological correspondence, spatial separation and training seem to be the most significant factors as evidenced by their effects on our CCE results and by previous observations^19,25^.

CCE scores were affected by training duration; spatial separation only affected CCE score in the extended-training subjects (Fig. 4). It seems that the short-training subjects did not have enough exposure to the feedback to reach a maximum level of incorporation, a qualitative finding observed elsewhere although on different timescales^26^. Therefore, to generate the physiological correspondence quantification model we used only extended-training data. The extended-training group had 80 minutes of practice with the feedback system, which is less training than would be typical for patients using this assessment. A patient with a novel feedback system will often complete a take-home trial, wearing the device for days to weeks, before running this assessment. Therefore, we consider only extended-training data and define the 80-minute duration as the minimum exposure necessary for this assessment to be effective. We expect the effect of training to plateau and that the model should be applicable to longer training times; nevertheless, this should be verified with an additional study.

Although the model’s goodness-of-fit seems low (R^2^=0.27), this is a consequence of the noisiness of human movement data and it can still be used to assess PNI feedback quality. However, the variability of CCE scores across individuals may make one-to-one comparison between individuals difficult. We based our model development on a population level analysis that, while statistically valid, could lead to misinterpretation of a single CCE result. Therefore, we recommend comparisons of CCE scores from the same individual across different feedback modalities or with different training periods. Alternatively, when data from several subjects are available, a population-level comparison can be made. A power analysis should be run to determine the number of subjects necessary to detect a certain level of improvement supplied by a novel feedback system^27^. To detect the maximum intergroup difference observed in our study (93.06ms), and assuming the observed overall variability across all 60 subjects (SD = 45.57ms, normalized to the mean within each group), three subjects would be needed to achieve a statistical power of 0.8 at a confidence level of 95%. Increasing the number of subjects tested would allow for smaller CCE score differences to be detected. For example, to detect half of the maximum difference observed (δ=46.53ms), eight subjects should be tested. The exact number of subjects required depends on the CCE score variability that will vary depending on the feedback system and patient population. In either the repeat-individual testing approach or the small-group population analysis, a washout period between repeat assessments is required to reduce a previously reported crossmodal congruency task learning effect^23^.

The low CCE score observed for skin deformation feedback (Fig. 3) was an unexpected result. Originally we hypothesized that using skin deformation feedback to represent the grasp force of the training movements would result in the highest level of incorporation. Skin deformation more closely resembles the physical activation of a grasping force compared to vibration and electrical stimulation. However, skin deformation feedback resulted in the lowest CCE score compared to electrical stimulation and vibration, a statistically significant result that does not appear to be the result of noise or random fluctuation. We confirmed that the poor incorporation of skin deformation was not due to mediocre actuation as latency results were consistent across feedback modalities. In some subjects the tactor may have been in contact with the skin at a zero force level. Variable skin contact would result in variable perception across subjects as the discrete initial skin contact has been shown to be important in improving feedback effectiveness^28^. The low incorporation of skin deformation feedback could alternatively be explained by long-term depression of afferents due to repeated stimulation or slipping actuators; both explanations would be supported by an observed change in detection threshold over the course of the experiment (see Supplementary Data S1). A future study is planned to combine the CCE score assessment with an outcome metric that assesses feedback uncertainty to more carefully characterize the utility of the skin deformation feedback^29^.

Subjects performed very well using skin deformation feedback (Supplementary Fig. S2), but we still observed poor incorporation. The inversely proportional trend between motor performance and CCE score may seem counterintuitive but implies that performance and incorporation are distinct, or even competing concepts. An individual’s quantifiable motor performance may be inflated through the adoption of alternative strategies and compensatory movements^30,31^. Further, motor performance does not necessarily correspond to other important aspects of prosthesis use such as device acceptance^13^, phantom pain reduction^14,15^ or cognitive burdens^16^. Therefore, clinical movement assessments relying only on motor performance may not be suitable to analyze the performance of PNIs. These assessments also suffer from other limitations such as a reliance on movement timing and variability introduced by rater subjectivity^32^. Available motor assessments may be sufficient to monitor clinical progress but no single outcome metric captures all relevant performance information^33^. The CCE-based assessment and supporting model we have presented could augment the battery of performance-based assessments currently in use to provide more detailed insight into PNI performance.

We have presented a CCE score assessment coupled with a data-driven model that can quantify the physiological correspondence of a feedback modality. This approach represents a way to provide more informative assessment of prosthetic feedback systems. Further steps will require the clinical validation of this assessment tool in patient populations, such as amputees outfitted with peripheral nerve feedback systems. Novel feedback systems for amputees require novel assessment tools; this work provides an advanced outcome metric to fill that need.

## Methods

### Participant recruitment

Participants were recruited by word-of-mouth and provided informed consent under the guidelines and approval of University of New Brunswick’s Research Ethics Board. Sixty volunteer participants completed the study [mean age = 31.9yrs, range = 18 – 76yrs, 22 female, 5 left-handed]. Participants were randomly assigned to a treatment condition which specified feedback modality [vibration, electrical stimulation or skin deformation], training duration [short or extended], and spatial separation between visual and experimental feedback [low or high].

### Bypass prosthesis

Subjects first trained using a bypass prosthesis with myoelectric control and embedded force sensors in the thumb and index finger that proportionally drove experimental feedback. The bypass is described in detail elsewhere^24^.

Skin deformation feedback was applied using linear mechanotactile haptic tactors attached to the subject (design courtesy of the University of Alberta^34^). Each tactor used a rack and pinion gear system to convert rotational motion generated by a servo motor (HiTec, HS-35HD) to linear motion that was applied to the subject’s skin via an 8mm diameter domed head. Measured force from the sensorized prosthetic hand (custom retrofitting of Ottobock MyoHand VariPlus Speed by HDT Global) was mapped to servo displacement. Zero force was mapped to a displacement that was a step below the minimum detectable level. The maximum displacement was based on the current draw of the servos and limited to approximately 100mA. This level was selected to keep the actuation at a level below which the plastic rack and pinion system would not slip. During the training phase, the tactor displacements were proportionally controlled to match the measured forces on the thumb and index finger of the instrumented prosthetic hand. During the CCE score assessment, the tactors were displaced to approximately 20-25% of the maximum experienced during the training phase.

Vibration feedback was provided by two 10mm linear resonant actuators (LRAs: Precision Microdrives, C10-100) taped to the skin with medical tape (3M, Micropore). During the training phase, the LRAs were proportionally controlled to correspond to the measured forces on the thumb and index finger of the instrumented prosthetic hand. During the CCE score assessment, the stimuli were set to approximately 20-25% of the maximum intensity experienced during the training phase.

Electrical stimulation was provided by a 2-channel TENS electro-stimulator (Proactive, Pulse). The device was modified such that the electrical stimulation intensity could be controlled with isolated analog outputs from a myRIO embedded hardware system (National Instruments). During the training phase, the stimulator outputs were proportionally controlled to produce paresthetic sensations that corresponded to the measured forces on the thumb and index finger of the instrumented prosthetic hand. During CCE score assessment, the stimuli were set to the maximum intensity experienced during the training phase. The protocol for electrical stimulation was modified compared to the other modalities to limit participant discomfort and avoid painful percepts.

For feedback with low spatial separation, the experimental feedback was applied at the fingertip to match the visually-observed contact point on the prosthetic hand. For feedback with high spatial separation, the actuators were attached to the wrist for skin deformation and vibration. For the electrical stimulation low spatial separation group, the self-adhesive electrical stimulation pads were wrapped around the index finger or thumb. Electrical stimulation on the wrist interfered with EMG control signals so for the high spatial separation group the pads were placed on the back of the hand near the major knuckles of the index finger and thumb.

Feedback detection thresholds were measured for each subject to calibrate the stimulation before the training phase. The stimulus intensity was slowly increased until the subject indicated that the stimulation was felt. This was repeated three times and the lowest reported stimulus level was used to set the range of stimulus. A proportional mapping was used to convert the hand’s force detection range to the subject’s stimulus detection range. The low end of the force detection range was set slightly below the reported detection threshold (~1% PWM duty cycle decrease for vibration and skin deformation feedback; ~10mV decrease for electrical stimulation). The maximum feedback was set to correspond to 1.2x the breaking threshold of the heaviest egg (19.4N). The maximum stimulus level was set based on the type of feedback. The LRAs were set to their maximum achievable intensity for the maximum stimulus level. The electrical stimulus maximum level was set based on the subject’s comfort and to avoid muscle twitch. The feedback detection threshold was measured again after the training phase, immediately preceding CCE score assessment.

The one-degree-of-freedom prosthetic hand of the bypass was controlled with a Complete Control (Coapt) pattern recognition system. Subjects trained hand open and close control using isometric wrist flexor and wrist extensor muscle contractions using the commercial software provided by Coapt.

### Training

Subjects in the short-training group completed five training sessions, each lasting ≤10 minutes with 10-minute intervening breaks, for 50 minutes of total training. Extended-training subjects completed eight sessions for 80 minutes of total training. In each training session subjects attempted to move instrumented mechanical eggs of three different weights and “breaking” thresholds over a 5cm high barrier. The lightest egg weighed 2.78N with a breaking threshold of 6.84N. The medium-weight egg weighed 5.45N with a breaking threshold of 10.52N. The heaviest egg weighed 9.55N with a breaking threshold of 16.19N. Each session ended after 100 movement attempts or ten minutes, whichever occurred first. Successful and unsuccessful movements were recorded manually by the experimenter. When too much force was applied to the mechanical egg, an on-egg LED would illuminate to indicate a broken egg. After breaking a mechanical egg, the subject had to release the egg and restart the movement. Subjects wore earplugs and over-ear noise-canceling headphones playing Brownian noise to mask actuator and background noise.

### CCE assessment

The CCE score assessment and associated hardware is as described in Gill et al.^23^. Subjects completed three familiarization sessions of ten trials each and then four assessment blocks of 64 trials each. Subjects were seated beside a height adjustable table that was set to a comfortable height. A pillow was placed under each subject’s arm to ensure vibrations were not transmitted through the table surface. CCE score for each block was computed as mean congruent time minus mean congruent time. The overall CCE score was calculated as the median of the scores from the four blocks.

### Statistical analysis

Statistical analysis was run using IBM’s SPSS Statistics and MATLAB software. A multi-way ANOVA was run with dummy categorical variables used to represent feedback modality, spatial separation and training level. Effect sizes were calculated as ω2 and reported as the square root, ω.35 For the linear regression analysis (see equation (1)), only extended-training data were used. CCE score was the dependent variable and Feedback Location and Physiological Correspondence were the independent variables. The Feedback Location variable was set to either zero (distances of 0 to 3 cm, at or near the fingertips) or one (distances of more than 12 cm from finger tips, on the wrist or back of hand). Physiological correspondence was set as the mean CCE score for a particular feedback type (71 for skin deformation feedback, 120.5 for vibration, 84.1 for electrical stimulation).

### Data availability

All data are available in the Supplementary Data S2 file that accompanies this manuscript.

## Acknowledgements

The authors thank Dr. Satinder Gill, Wendy Hill, and Heather Daley for their assistance in the implementation of this work. This work was partially funded by the Defense Advanced Research Projects Agency (DARPA) through the HAPTIX program (award number N66001-15-C-4015) and the New Brunswick Health Research Foundation (NBHRF) through an Establishment Grant.

## Author contributions

DB wrote the main manuscript text and prepared the figures. AW designed and built the experimental hardware and collected all data. JS edited the manuscript. All authors contributed to the experimental design, data analysis and manuscript review.

## Additional information

### Competing financial interests

J. Sensinger is a co-owner of Coapt Engineering whose product was used in this study.

## Supplementary information

**Supplementary Table S1.** Experimental conditions for 60 able-bodied subjects. Spatial separation is marked as Ø for 3cm or less and ✓ for greater than 12cm.

| Feedback modality → | Vibration |  |  |  | Electrical stimulation |  |  |  | Skin deformation |  |  |  |
| --- | --- | --- | --- | --- | --- | --- | --- | --- | --- | --- | --- | --- |
| Training level → | Short |  | Extended |  | Short |  | Extended |  | Short |  | Extended |  |
| Spatial separation → | ∅ | ✓ | ∅ | ✓ | ∅ | ✓ | ∅ | ✓ | ∅ | ✓ | ∅ | ✓ |
| # of subjects → | 5 | 5 | 5 | 5 | 5 | 5 | 5 | 5 | 5 | 5 | 5 | 5 |

**Supplementary Figure S1.**
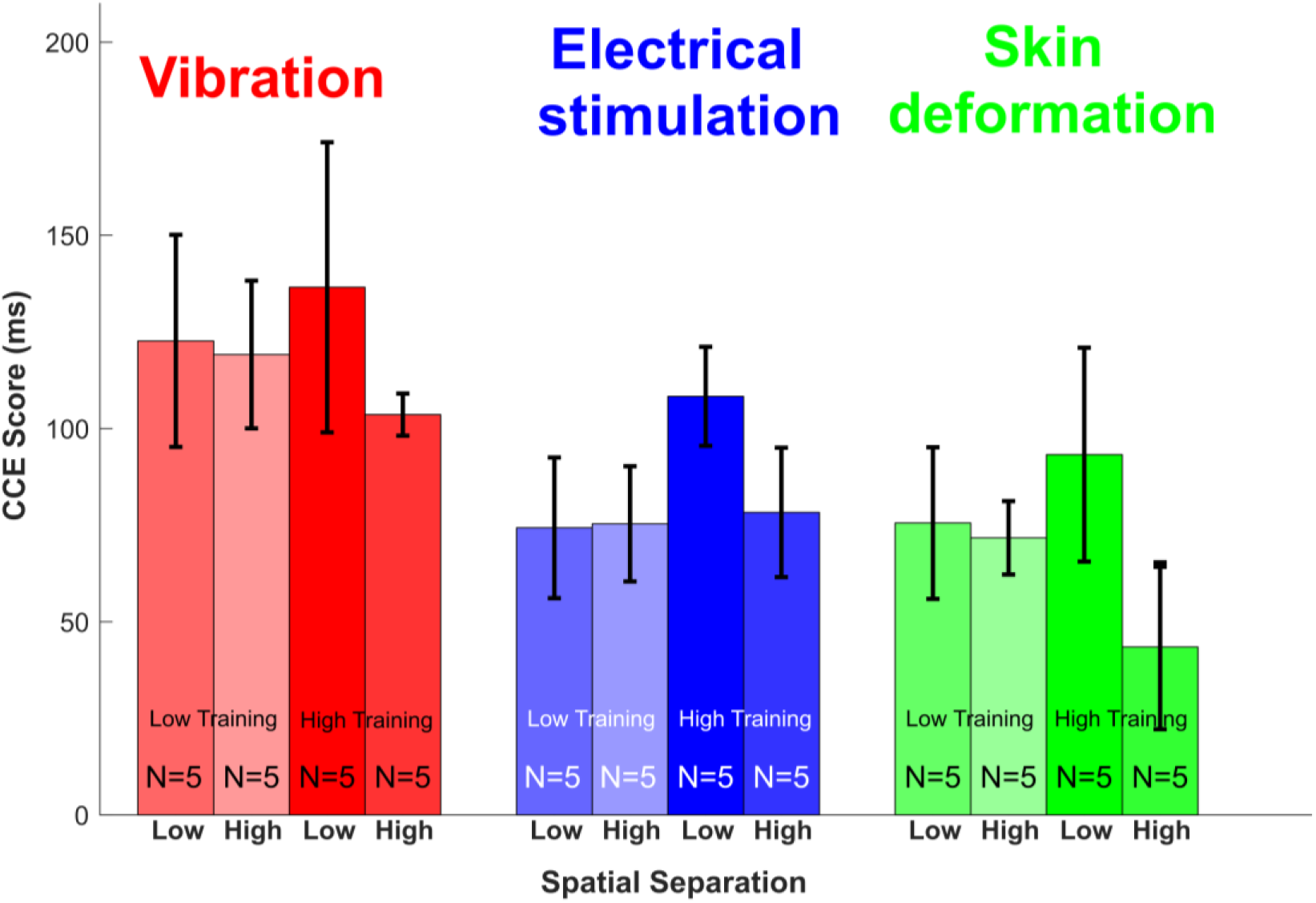
*Mean CCE scores and standard error from all 12 treatment groups.*

### Supplementary Data S1. Additional investigation of low CCE score with skin deformation feedback

Surprisingly, skin deformation feedback resulted in the lowest CCE score, corresponding to the lowest level of incorporation (Fig. 3). We sought to investigate this result by analyzing the success rate of movements during training. CCE score tended to increase as the percent of successful egg movements (no drops and no breaks) decreased (Supplementary Fig. 2). Differences in success rates between feedback modality were not significantly different determined with one-way ANOVA (F(2,57)=2.48, p = 0.093).

The incorporation of skin deformation feedback may have been affected by the intensity, timing or other characteristics of the feedback provided. We observed a significant difference (unpaired t-test, p<0.05) in the change in detection threshold over the course of training between vibration (0% change) and skin deformation modalities (+51.4% average change). Detection thresholds were measured at the start of the training phase and at the end of training just before CCE score assessment. The detection threshold of the electrical stimulation feedback was set differently to avoid painful sensations and was not included in this analysis. There were no differences measured between the timing precision of the different feedback modalities. The initial position of the tactor may have affected the effectiveness of the skin deformation feedback. In some subjects the tactor may have been in contact with the skin or arm hair before any sensed force. In future studies, body hair should be shaved and the tactor should be initially positioned to ensure no contact with the subject at zero force levels.

**Supplementary Figure S2.**
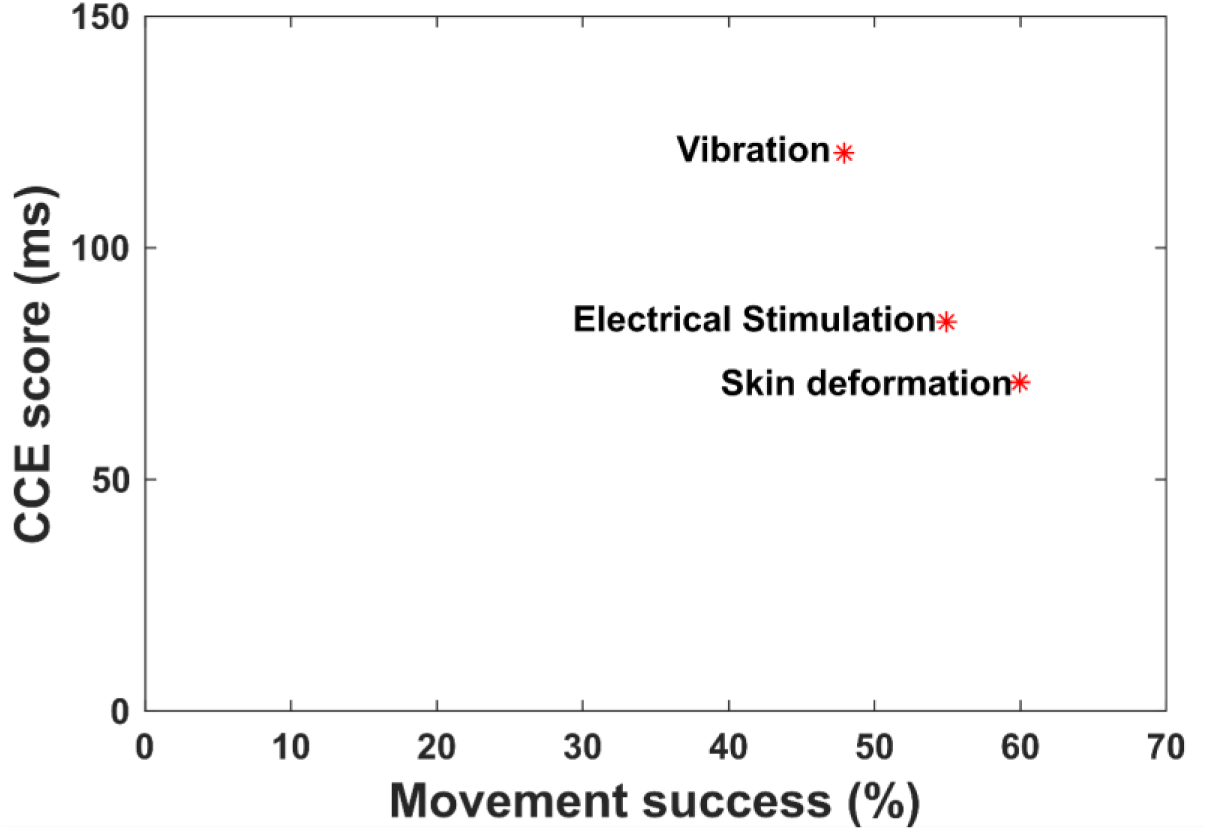
CCE score and movement success rate show an inversely proportional trend. Differences in movement success rates were not significantly different (one-way ANOVA; F(2,57)=2.48, p = 0.093).

## References

1. Østlie, K. et al. Prosthesis rejection in acquired major upper-limb amputees: a population-based survey. Disabil. Rehabil. Assist. Technol. 7, 294–303 (2011).

2. Biddiss, E. A. & Chau, T. T. Upper limb prosthesis use and abandonment: A survey of the last 25 years. J. Prosthet. Orthot. Int. 31, 236–257 (2007).

3. Raspopovic, S. et al. Restoring natural sensory feedback in real-time bidirectional hand prostheses. Sci. Transl. Med. 6, 222ra19 (2014).

4. Tan, D. W. et al. A neural interface provides long-term stable natural touch perception. Sci. Transl. Med. 6, 257ra138 (2014).

5. Ortiz-Catalan, M., Håkansson, B. & Brånemark, R. An osseointegrated human-machine gateway for long-term sensory feedback and motor control of artificial limbs. Sci. Transl. Med. 6, 257re6 (2014).

6. Clark, G. A., Ledbetter, N. M., Warren, D. J. & Harrison, R. R. Recording sensory and motor information from peripheral nerves with Utah Slanted Electrode Arrays. Conf. Proc. IEEE Eng. Med. Biol. Soc. 2011, 4641–4644 (2011).

7. Mastinu, E., Doguet, P., Botquin, Y., Håkansson, B. & Ortiz-Catalan, M. Embedded System for Prosthetic Control Using Implanted Neuromuscular Interfaces Accessed Via an Osseointegrated Implant. IEEE Trans. Biomed. Circuits Syst. 11, 867–877 (2017).

8. Christie, B. P. et al. Long-term stability of stimulating spiral nerve cuff electrodes on human peripheral nerves. J. Neuroeng. Rehabil. 14, 70 (2017).

9. Mathiowetz, V., Volland, G., Kashman, N. & Weber, K. Adult norms for the Box and Block Test of manual dexterity. Am. J. Occup. Ther. 39, 386–391 (1985).

10. Mathiowetz, V., Weber, K., Kashman, N. & Volland, G. Adult Norms for the Nine Hole Peg Test of Finger Dexterity. Occup. Ther. J. Res. 5, 24–38 (1985).

11. Light, C. M., Chappell, P. H. & Kyberd, P. J. Establishing a standardized clinical assessment tool of pathologic and prosthetic hand function: Normative data, reliability, and validity. Arch. Phys. Med. Rehabil. 83, 776–783 (2002).

12. Hermansson, L. M., Fisher, A. G., Bernspång, B. & Eliasson, A.-C. Assessment of capacity for myoelectric control: a new Rasch-built measure of prosthetic hand control. J. Rehabil. Med. 37, 166–171 (2005).

13. Marasco, P. D., Kim, K., Colgate, J. E., Peshkin, M. A. & Kuiken, T. A. Robotic touch shifts perception of embodiment to a prosthesis in targeted reinnervation amputees. Brain 134, 747–758 (2011).

14. Tyler, D. J. Neural interfaces for somatosensory feedback. Curr. Opin. Neurol. 28, 574–581 (2015).

15. Ortiz-Catalan, M. et al. Phantom motor execution facilitated by machine learning and augmented reality as treatment for phantom limb pain: a single group, clinical trial in patients with chronic intractable phantom limb pain. Lancet 388, 2885–2894 (2016).

16. Makin, T. R., de Vignemont, F. & Faisal, A. A. Neurocognitive barriers to the embodiment of technology. Nat. Biomed. Eng. 1, 0014 (2017).

17. Davis, T. S. et al. Restoring motor control and sensory feedback in people with upper extremity amputations using arrays of 96 microelectrodes implanted in the median and ulnar nerves. J. Neural Eng. 13, 036001 (2016).

18. Shannon, G. F. A comparison of alternative means of providing sensory feedback on upper limb prostheses. Med. Biol. Eng. Comput. 14, 289–294 (1976).

19. Spence, C., Pavani, F. & Driver, J. Spatial constraints on visual-tactile cross-modal distractor congruency effects. Cogn. Affect. Behav. Neurosci. 4, 148–169 (2004).

20. Zopf, R., Savage, G. & Williams, M. A. The Crossmodal Congruency Task as a Means to Obtain an Objective Behavioral Measure in the Rubber Hand Illusion Paradigm. J. Vis. Exp. 77, e50530 (2013).

21. Maravita, A., Spence, C., Kennett, S. & Driver, J. Tool-use changes multimodal spatial interactions between vision and touch in normal humans. Cognition 83, B25–34 (2002).

22. Frings, C. & Spence, C. Crossmodal congruency effects based on stimulus identity. Brain Res. 1354, 113–122 (2010).

23. Gill, S., Wilson, A. W., Blustein, D. & Sensinger, J. Crossmodal congruency effect as a metric of tool incorporation: assessment of metric attenuation caused by overexposure, and modifications to reduce attenuation. Preprint at https://www.biorxiv.org/content/early/2017/09/10/186825 (2017).

24. Wilson, A. W., Blustein, D. H. & Sensinger, J. W. A third arm - Design of a bypass prosthesis enabling incorporation. IEEE Int. Conf. Rehabil. Robot. 2017, 1381–1386 (2017).

25. Mayer, A. R., Franco, A. R., Canive, J. & Harrington, D. L. The effects of stimulus modality and frequency of stimulus presentation on cross-modal distraction. Cereb. Cortex 19, 993–1007 (2009).

26. Marini, F. et al. Crossmodal representation of a functional robotic hand arises after extensive training in healthy participants. Neuropsychologia 53, 178–186 (2014).

27. Lieber, R. L. Statistical significance and statistical power in hypothesis testing. J. Orthop. Res. 8, 304–309 (1990).

28. Cipriani, C., Segil, J. L., Clemente, F., ff Weir, R. F. & Edin, B. Humans can integrate feedback of discrete events in their sensorimotor control of a robotic hand. Exp. Brain Res. 232, 3421–3429 (2014).

29. Blustein, D. & Sensinger, J. W. Extending a Bayesian estimation approach to model human movements. Program No. 486.10. 2016 Neuroscience Meeting Planner. San Diego, CA: Soc. for Neurosci. (2016).

30. Hussaini, A., Zinck, A. & Kyberd, P. Categorization of compensatory motions in transradial myoelectric prosthesis users. J. Prosthet. Orthot. Int. 41, 286–293 (2016).

31. Hebert, J. S. & Lewicke, J. Case report of modified Box and Blocks test with motion capture to measure prosthetic function. J. Rehabil. Res. Dev. 49, 1163–1174 (2012).

32. Hermansson, L. M., Bodin, L. & Eliasson, A.-C. Intra- and Inter-Rater Reliability of the Assessment of Capacity for Myoelectric Control. J. Rehabil. Med. 38, 118–123 (2006).

33. Wright, V. Prosthetic Outcome Measures for Use With Upper Limb Amputees: A Systematic Review of the Peer-Reviewed Literature, 1970 to 2009. J. Prosthet. Orthot. 21, P3–P63 (2009).

34. Schoepp, K., Dawson, M., Carey, J. & Hebert, J. Design and integration of an inexpensive wearable tactor system. Myoelectric Controls Sympos. Fredericton, NB, Canada. **ID# 89**, (2017).

35. Field, A. Discovering Statistics Using SPSS. Los Angeles, CA.: SAGE Publications. (2009).

